# No Evidence for Long-range Male Sex Pheromones in Two Malaria Mosquitoes

**DOI:** 10.1101/2020.07.05.187542

**Authors:** Serge Bèwadéyir Poda, Bruno Buatois, Benoit Lapeyre, Laurent Dormont, Abdoulaye Diabaté, Olivier Gnankiné, Roch K. Dabiré, Olivier Roux

## Abstract

Cues involved in mate seeking and recognition prevent hybridization and can be involved in speciation processes. In malaria mosquitoes, females of the two sibling species *Anopheles gambiae s.s.* and *An. coluzzii* mate in monospecific male swarms and hybrids are rare. Long-range sex pheromones driving this behavior have been debated in literature but to date, no study has proven their existence or their absence. Here, we attempted to bring to light their existence. To put all the odds in our favor, we used different chemical ecology methods such as behavioral and electrophysiological assays as well chemical analyses, and we worked with mosquitoes at their optimal physiological mating state *i.e.* with swarming males during their natural swarming windows. Despite all our efforts, our results support the absence of long-range sex pheromones involved in swarm detection and recognition by females. We briefly discuss the implications of this finding in ecology, evolution and for control strategies.

## INTRODUCTION

Cues involved in mate seeking and mate recognition prevent hybridization and can be involved in speciation processes. In insects, these cues can be visual, acoustical or chemical with a short or a long-range action and are highly species specific (Alexander et al. 1997; Clements 1999).

In mosquitoes, many species mate in flight, within individual aggregations called “mating swarms” (Gibson 1945; Downes 1969; Savolainen 1978; Shelly and Whittier 1997; Sivinski and Petersson 1997). In malaria mosquitoes of the *Anopheles gambiae* complex, mating takes place at sunset in monospecific swarms containing a few males to thousands of them in which virgin conspecific females come to find a mate (Charlwood and Jones 1980; Marchand 1984; Charlwood et al. 2002; Diabaté et al. 2003, 2009; Howell and Knols 2009; Sawadogo et al. 2014). In West Africa, *An. coluzzii* and *An. gambiaes.s.* are often found in sympatry but they form distinct swarms spatially segregated and hybrids are rare (≈1%) (della Torre et al. 2001; Tripet et al. 2001; della Torre et al. 2005; Diabaté et al. 2006; Costantini et al. 2009; Diabaté et al. 2009; Sawadogo et al. 2013, 2014). No evidence for selection against hybrids was found and spermatheca analyses showed that these species mostly mate assortatively (Persiani et al. 1986; Tripet et al. 2001; Diabaté et al. 2005, 2007; Hahn et al. 2012; Pombi et al. 2017). This suggests that reproductive isolation between these two sibling species is achieved by strong pre-mating reproductive barriers (della Torre et al. 2001; Diabaté et al. 2007; Lehmann and Diabaté 2008). In addition, since females usually only mate once in their lives (Clements 1992), errors in the choice of mate should be costly and fall under negative selection. Consequently, one would expect to find specific cues that lead females to conspecific male swarms. However, the way females are attracted to swarms is unknown.

In species of the *An. gambiae* complex, several cues have been identified to play a role in bringing sexes together. Acoustic cues were shown to be involved in close-range recognition, the male and the female adjusting their respective wing-beat frequencies to converge on a shared harmonic frequency (Gibson et al. 2010; Pennetier et al. 2010). However, it was recently demonstrated that females of some *Anopheles* species are able to detect swarm sounds only in a very close vicinity. Thus, they are unable to use swarm sound to locate or identify swarms at long range (Feugère et al. 2021). Visual cues also play an important role in species segregation with *An. coluzzii* males swarming over contrasted visual ground markers (marker, hereafter) and *An. gambiae* males swarming over bare ground (Diabaté et al. 2009). However, a recent work showed that, like *An. coluzzii, An. gambiae* males also use visual markers but rather to locate their swarm at a distance from the marker (Poda et al. 2019). Moreover, females also use these markers to form swarms in the absence of males suggesting that females may use these markers to join the swarm location (Poda et al. 2019). Nevertheless, the distance between two heterospecific swarms using the same visual ground marker can be about 2 m in semi-field conditions (Poda et al. 2019) and it is still unknown from which distance mosquitoes can see such markers, suggesting that females could cross the location of heterospecific swarms by accident. Chemical cues have also been under investigation in *Anopheles* species. Heptacosane, a cuticular hydrocarbon, enhances the interaction between males and females (Wang et al. 2021). This compound, however, can only be perceived by contact and can consequently be involved only in mate recognition during courtship and to stimulate acceptance by females. Moreover, the absence of assortative mating in confined heterospecific males and females in laboratory cages or indoor swarms (Dao et al. 2008), suggests that close-range mating cues, if they exist, cannot ensure total reproductive isolation by themselves. Species isolation is thus likely to occur through long-range and specific swarm recognition cues acting as a first barrier in pre-mating processes which prevent hybridization by limiting contact between sexes of the different species.

To our knowledge, volatile sex pheromones in the *An. gambiae* complex have never been brought to light without ambiguity. Charlwood et al. (2002) reported an absence of response of *An. gambiae* males in natural swarms to squashed females on filter paper or to living females in a net cage. However, in natural conditions, females are the ones attracted to male aggregation sites. Consequently, males should be the ones that emit long-range pheromones and should be the attractive sex. Nevertheless, a laboratory study failed to demonstrate any attractiveness of dead males to virgin females in a Y-tube olfactometer (Gomulski 1988). However, the difficulties in highlighting the existence of such pheromones might be due to an emission in very low quantities and/or exclusively during swarming. Indeed, in sandfly species, which also mate in large aggregations of males, it was shown that the concentration of male-produced sex pheromones is greater in larger swarms, and hence, the chances for individual males of mating with a conspecific female are also greater (Kelly and Dye 1997; Bray et al. 2010). Similarly, Diabaté et al. (2011) showed that *An. coluzzii* females were more frequently attracted to large swarms, suggesting a potential additive effect of cues released by males. Males gathering in large swarms may increase their “detectability” in the female’s olfactory landscape (Shelly and Whittier 1997). In addition, in many insects, pheromone release and receptivity were shown to respond to diel rhythms or even strict time windows of only a few minutes or hours (Bjostad et al. 1980; Merlin et al. 2007; Rund et al. 2013; Levi-Zada et al. 2014) or even to be dependent of the presence of conspecifics (Andersson et al. 2007; Robledo and Arzuffi 2012). Nevertheless, a recent work by Mozūraitis et al. (2020) and concomitant to a first version of the present work (Poda et al. 2021), highlighted the existence of five volatile compounds potentially emitted by *Anopheles* males. According to the authors, these might be involved in aggregation behavior, attracting both males and females, and increasing the insemination rate. However, though an interesting biological activity was described by the authors, the claim that these compounds are male swarming aggregation pheromones must be considered with caution and it requires further investigations. Indeed, these five compounds (acetoin, sulcatone, octanal, nonanal and decanal) are very frequently found in nature and, as reported by the authors, have been shown to have a biological activity in *Anopheles* mosquitoes in very different contexts from the reproductive behavior. They were repeatedly found in human and animal body odor (Verhulst et al. 2010; Pandey and Kim 2011; Dormont et al. 2013b, a; Tchouassi et al. 2013; McBride et al. 2014), breath (Poli et al. 2010; Calenic and Amann 2014; Filipiak et al. 2014; Cainap et al. 2020), and in host-plants (Dekel et al. 2019). These represent the main sources of food (blood and sugar meals, respectively) for *Anopheles* mosquitoes. As a matter of fact, octanal, nonanal and decanal are part of a blend used in some traps to mimic mammalian host odor for several species of mosquito vectors (Tchouassi et al. 2013; Nyasembe et al. 2014). Some of these compounds were also found associated to oviposition sites (Suh et al. 2016; Wondwosen et al. 2016, 2018) and to ambient air (Kostiainen 1995; Kruza et al. 2017). In addition, sulcatone is thought to be responsible for discrimination between humans and animals in human-seeking mosquitoes (McBride et al. 2014). Thus, it is not surprising that they trigger a strong biological/flight activity in mosquitoes.

Nevertheless, the discovery of sex pheromones or a highly attractive blend in malaria mosquitoes would be a precious step toward the development of new control and monitoring strategies. Furthermore, sex or aggregation pheromones could help in designing sexually competitive mosquitoes in sterile insect and gene drive techniques. Here, we investigated the existence of long-range sex pheromones in *An. gambiae* and *An. coluzzii* mosquitoes, that may allow the females to detect, recognize and track conspecific male swarms. On the premise that such pheromones could be produced only by males during swarming windows and to put all the odds in our favor, we used different chemical ecology methods, always with living swarming mosquitoes during their natural swarming windows. First, we investigated the long-range behavioral response of females exposed to the volatile blend from male swarms in an olfactometer. Second, we collected and analyzed volatile organic compounds (VOCs) with different methods on both laboratory-induced swarms and natural swarms. And third, we tested for an antenna-electrophysiological response of females to male swarm VOCs. As much as possible, we used both recently colonized mosquitoes and large experimental set-ups to ensure males produced a free swarming behavior. In addition, we replicated the main experiment by Mozūraitis et al. (2020) to search for the five compounds. We specifically added two new controls, one to check if these compounds were male specific, and the other to discard a potential environmental/laboratory pollution.

## Methods and Materials

### Mosquitoes

Mosquitoes used in behavioral experiments and for both VOC collection and electrophysiological analyses were from colonies raised from wild gravid females collected in inhabited human dwellings in Burkina Faso (West Africa). *Anopheles coluzzii* were collected in 2017 in Bama and *An. gambiae* in 2015 in Soumousso. Bama is a village located in a rice-growing area located 30 km North of Bobo-Dioulasso (11°24’14”N; 04°24’42”W) where previous studies showed that the *Anopheles* population is almost exclusively composed of *An. coluzzii* (Mosqueira et al. 2015; Poda et al. 2018). Soumousso is a typical Guinean savannah village located 30 km North-East of Bobo-Dioulasso (11°00’46”N, 4°02’45”W) where *An. coluzzii* and *An. gambiae* coexist, with a predominance of the latter (Diabaté et al. 2004, 2006). Gravid females were placed individually in oviposition cups containing tap water. After oviposition, females were identified to species by routine PCR-RFLP (Santolamazza et al. 2008). The larvae were gathered according to their species and reared in tap water, fed with Tetramin® Baby Fish Food (Tetrawerke, Melle, Germany) *ad libitum.* Adult mosquitoes were held in 30 × 30 × 30 cm mesh-covered cages and provided with a 5% glucose solution *ad libitum.* Insectarium conditions were maintained at 27±2 °C, 70±10% RH and 12L:12D. The colonies were refreshed twice a year with F1 from mosquito females caught in the wild.

Mosquitoes used in the electrophysiological study were transferred as eggs from Burkina Faso to France. On their arrival, eggs were allowed to hatch in osmosed water and the larvae were fed with Tetramin® Baby Fish Food *ad libitum*. Adult mosquitoes were held in 20 × 20 × 20 cm mesh-covered cages and provided with honey diluted at 5% *ad libitum*. Females were fed with rabbit blood on a PS6 Power Unit (Hemotek, Blackburn, UK) for egg production. Mosquitoes were reared in a laboratory climate chamber KBF-S720 (BINDER Gmbh, Tuttlingen, Germany) at 27±2 °C, 70±10% RH and 12L:12D with a sunset time programmed at 3:00pm (synchronization of swarming/mating time and electrophysiological test time to ensure an optimal receptivity of females). Mosquitoes used for the experiments were sexed early after emergence and sexes were kept in separate rearing cages to prevent mating.

Mosquitoes used to replicate the experiment by Mozūraitis et al. (2020) were from a 15-year-old colony of *An. gambiae* (Kisumu strain) reared in the IRD laboratory of Montpellier. They were maintained at 27 ±2 °C, 80 ± 10% RH, with a photoperiod cycle of 12h light: 12h dark and reared as described above.

### Long-range behavioral response of virgin females to swarm volatile organic compounds

#### Olfactometer setup

Bioassays were conducted in a dual-port olfactometer originally designed to study host preference in the *An. gambiae* complex (Lefèvre et al. 2009, 2010; Vantaux et al. 2015; Nguyen et al. 2017). The odor source container was modified to enclose male swarms in boxes made of transparent plexiglass (L × W × H: 60 × 60 × 120 cm; “swarming boxes” hereafter). Each swarming box was connected by a PVC air vent hoses (L × Ø: 600 × 10 cm, W3-65014-HQ4, HQ, USA) to a collecting glass box (L × W × H: 32 × 33.5 × 44 cm) which was linked to a mesh-covered releasing cage (L × W × H: 50 × 40 × 40 cm) by a glass tube (L × Ø: 60 × 10 cm) (Fig. 1). A custom-made electric fan was located at the mid-length of each air vent hose and drew air from the swarming boxes (odor-sources) to the releasing cage, providing an odor-laden air current against which mosquitoes in the releasing cage were induced to fly. The air flow was controlled thanks to two mechanisms; first, a power regulator (HQ-Power PS1502A, Velleman, Gavère, Belgium) connected to fans; and second, iris dampers (10 cm Ø; CIR D100, France air, France) connecting the air vent hoses to the collecting boxes. The openings of the air vent hoses on both the swarming and collecting boxes sides were covered with nets to prevent mosquitoes from flying into the air vent hoses. The swarming boxes (odor-sources) were located side-by-side outdoors and the olfactometer inside a room (Fig. 1). The air speed in the releasing cage was regulated at 18±2 cm.s^-1^ using an anemometer (Model 425, Testo, Forbach, France) and the room temperature and relative humidity were set at 27±2 °C and 80±10% RH thanks to an air conditioner and a humidifier (Defensor 505, Condair, Croissy-Beaubourg, France), respectively.

**Fig. 1:**
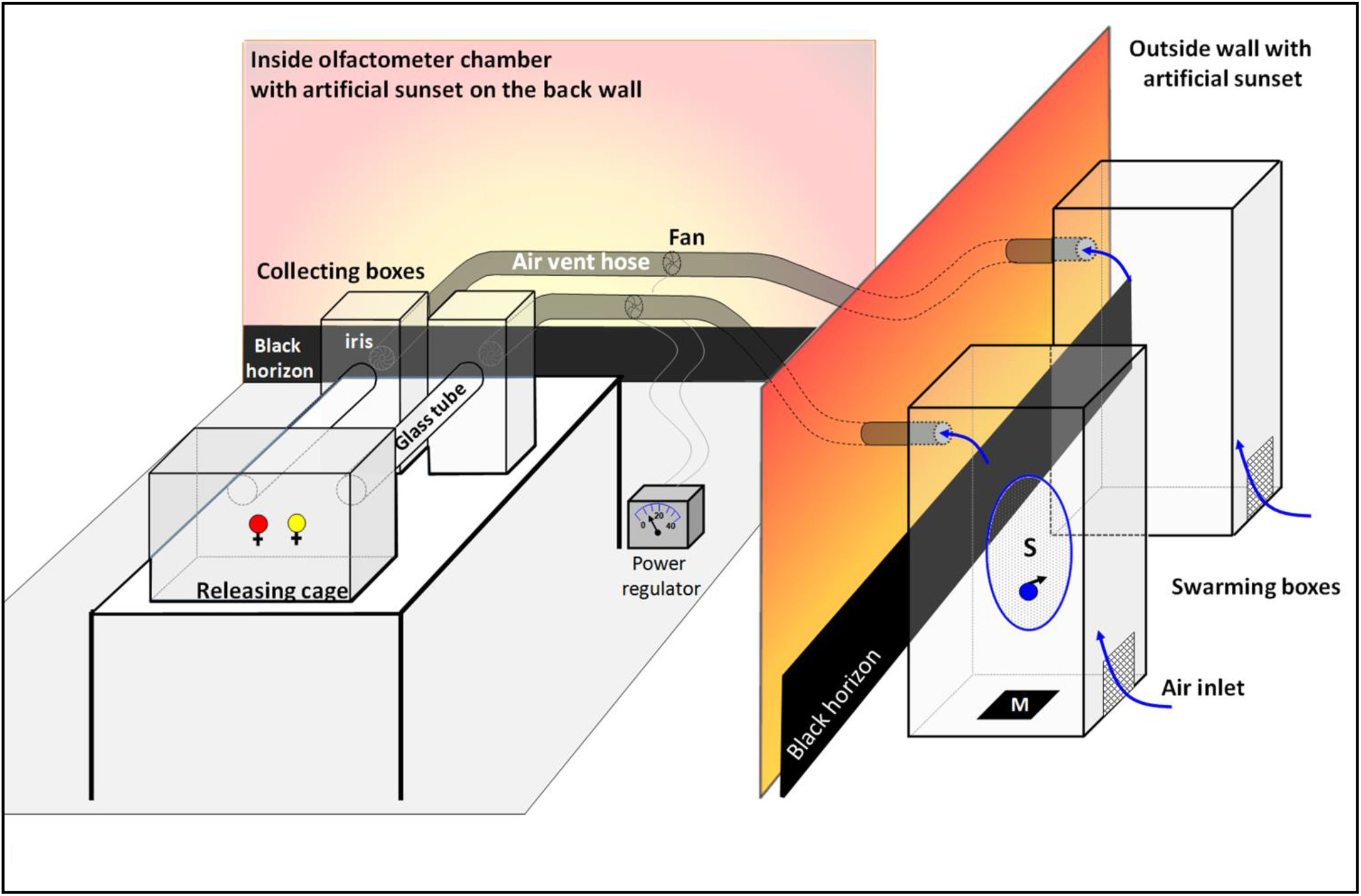
Schematic of the olfactometer set up. S: swarm; M: marker. Not at scale.

#### Bioassays

##### Test preparation

In the morning, about 8 hours before the time of the test, 100 2-to 3-day-old virgin females of *An. coluzzii* and *An. gambiae* were colored with two different colored powders (Luminous Powder Kit, BioQuip, Rancho Dominguez, California, USA) according to the species. In addition, about 500 4-to 5-day-old *An. coluzzii* or *An. gambiae* males were released into a swarming box to allow them to acclimate. Both males and females were provided with a 10% glucose solution and were kept under insectary conditions until the time of the test. To avoid biases during the test due to both humidity and odor as a result of manipulation or bacterial proliferation, the empty swarming box (control box) was also provided with glucose.

##### Test execution

Bioassays were carried out at sunset, when males naturally swarm. About 30 min before sunset time, the swarming boxes were moved outside, the cups containing the glucose pads were removed and visual markers made of a 20 × 20 cm black cloth were placed either under or next to the boxes for *An. coluzzii* or *An. gambiae,* respectively (see Poda et al. 2019). These markers allowed to trigger swarming behavior and to stabilize swarms in the middle of the box. For the same purpose, we provided an artificial twilight horizon made of a 40 W incandescent bulb (2,500 K) located on the floor between a white wall and a 50 cm high black horizon (Facchinelli et al. 2015; Niang et al. 2019; Poda et al. 2019). Inside, about 10 min before introducing females, the fans were switched on in order to purge the air vent hoses and the air flow was set at 18±2 cm.s^-1^. The glass tube openings on the releasing cage side were covered with nets to prevent female mosquitoes from flying into the collecting boxes before the test began. The complete olfactometer, except the end of the releasing cage, was covered with a white cloth to eliminate visual bias during the test and to provide a diffused light. Sunset light was allowed to enter the olfactometer room until dark and an additional artificial twilight made of a 40 W incandescent bulb (2,500K) projected on the wall of the room on the side of the collecting boxes was also provided. Thirty minutes before sunset, about 100 females of each species (N=200, 1 replicate) were released simultaneously into the releasing cage of the olfactometer allowing them to acclimate.

When the males started to swarm *(i.e.* flying in loops at a constant location within the swarming boxes, see Downes 1969; Poda et al. 2019), the nets covering the glass tube openings and preventing access of females to the collecting boxes were removed thanks to threads that were attached to the nets. The females were allowed to respond for 20 min (swarm duration), then the glass tube openings were covered again with nets and females that reached the upwind collecting boxes were removed with a mouth aspirator, killed by freezing and identified according to their coloration (species) under a binocular (LEICA S6E) and counted. The remaining mosquitoes inside the releasing cage were also removed. After each test (1 per day), the olfactometer was cleaned with 95% ethanol to remove odor contaminants. All materials were handled with gloves to avoid contamination with skin compounds.

##### Test combinations

Four combinations of choice tests were performed: i) *An. coluzzii* male swarm *vs.* empty box (hereafter, *An. coluzzii* test); ii) *An. gambiae* male swarm *vs.* empty box (hereafter, *An. gambiae* test), iii) *An. coluzzii vs. An. gambiae* male swarms (hereafter, *An. coluzzii vs. An. coluzzii* test) and iv) empty *vs.* empty box (hereafter, control test). We ran 12 replicates per combination and females were used only once. Males were used twice in two consecutive days. In that case, a 10% glucose solution was introduced into the two swarming boxes after the test and boxes were kept under insectarium conditions until the next day. The swarming boxes were cleaned with ethanol between each batch of males. To avoid biases, the matching of species and colors was switched between each test, and we alternated the treatments (mosquitoes or control) between the swarming boxes and the right and left arm of the olfactometer. We assessed the instrumental and arm bias through an empty *vs.* empty box choice test (control test). The olfactometer did not present any symmetrical biases (Fig. S1).

### Collection of swarm volatile organic compounds (VOCs)

Because compounds can be emitted at a given time and only produced in tiny quantities at individual scale, we chose to work directly on swarms. Three different sampling methods were used; two methods were run in a laboratory setup (Niang et al. 2019) and one in the field.

#### Dynamic headspace collection in the laboratory

##### VOC collection setup

The collection of swarm VOCs was performed in a room specially designed to stimulate *Anopheles* swarming behavior thanks to a set of visual stimuli (see Niang et al. (2019) for details). The headspace volume consisted of a 50 × 50 × 50 cm transparent plexiglass box (extraction box). Such a volume was a trade-off between a volume large enough to allow males to display a swarming behavior without too much constraints and a volume small enough to be sucked up in a reasonable time. Entrance and exit flow rates were maintained by two pumps and regulated by flowmeters. The entrance flow was higher than the exit flow ensuring that the system was continuously purged to compensate for the inevitable leaks, and that no contaminated outside air would enter the system. At the entrance side, ambient air was purified in a glass charcoal filter and then humidified passing through deionized water. Exit flow passed through an odor trap to adsorb VOCs. Three different odor traps were used for the chemical analyses and electrophysiological analyses. Tenax-TA/Carbotrap sorbent stainless steel 6mm diameter tubes (Gerstell, Mülheim, Germany) and Porapak-Q sorbent VCT glass tubes (ARS, Gainesville, Florida, USA) were used for chemical analyses. Porapak-Q sorbent VCT tubes and home-made sorbent Micro-traps were used for electrophysiological analysis. Micro-traps were constituted by ChromatoProbe® quartz microvials of Varian Inc. (length: 15 mm; inner diameter: 2 mm), previously cut closed-end and filled with 3 mg of a 1:1 mix of Tenax-TA and Carbotrap® (60–80 and 20–40 mesh, respectively; Sigma Aldrich, Munich, Germany)(See Dormont et al. 2013b). Flow rates and VOC collection durations were dependent on the odor trap (See Table 1).

**Table 1:**
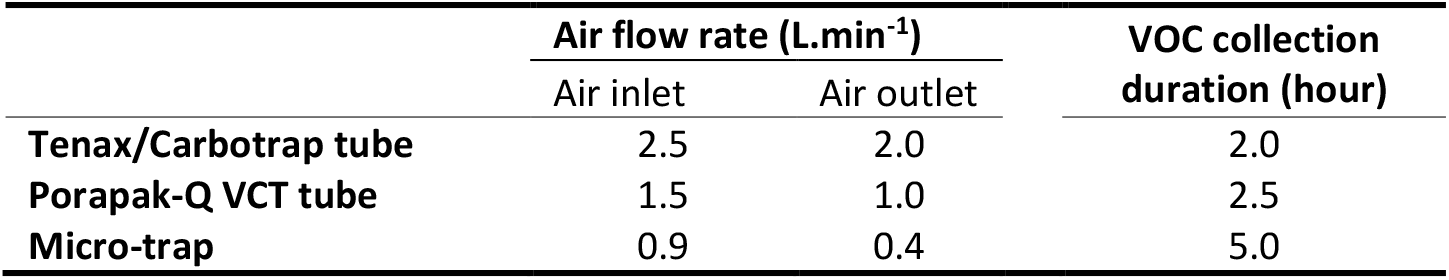
Air inlet and outlet rates, and VOC collection duration according to the type of trap used.

##### VOC collection

About 500 4- to 6-day-old males of *An. gambiae* or *An. coluzzii* were transferred into the extraction box about 30 min before swarming time. Clean air was pushed into the box at a rate of 10 L.min^-1^ to purge the box from both human and mosquito container odors before connecting it to the odor trap. Five minutes after males started to swarm *(i.e.* flying in loops at a constant location within the box), the air flow was adjusted at the required rate and the odor trap connected. Forty minutes later, swarming behavior was stopped by turning off the light and the VOC collection continued until the required collection time was reached (see Table 1). After VOC collection, samples were stored at 4 °C until analysis. After males were removed, the box was cleaned with 95% ethanol and then flushed with clean air for at least 5 hours to remove the odor contaminants left from the previous extraction. A total of 30 swarm extracts were collected; 17 *An. coluzzii* swarms (5, 8 and 4 with Tenax/Carbotrap tubes, Porapak-Q VCT tubes and Micro-traps, respectively) and 13 *An. gambiae* swarms (6 and 7 with Tenax/Carbotrap tubes, and Porapak-Q VCT tubes, respectively). Control consisted of a clean empty box. Before they were sent and used in Burkina Faso, adsorbents were cleaned as follows. Tenax-TA/Carbotrap tubes were heated at 250 °C for 30 min under a 30 mL.min^-1^ flow then sealed. Porapak-Q VCT tubes were eluted with 3 mL of hexane then packed individually in a nalophan bag. Chromatoprobes were heated for 2 h at 270 °C by 100 chromatoprobes under a 100 mL.min^-1^ nitrogen flow, then packed in a glass vial with a Teflon cap.

#### VOC collection from natural swarms

VOCs were collected from a natural swarm of *An. coluzzii* in the village of Bama. Static sorptive collections of volatiles from a non-enclosed swarm were performed with Twisters® (100 μm PDMS stir bar; Gerstel, Mülheim, Germany; Bicchi et al. 2000; Tienpont et al. 2000). The swarm flew about 2.5 to 3.5 m above the ground. Thus, we used a 1.5 m long glass stick (Glaswarenfabrik Karl Hecht GmbH & Co, Germany) with a metallic push pin head glued at the tip and covered with nalophan to introduce the magnetic twisters directly into the swarm. Control twisters were placed about 3 m away upwind from the swarm. VOC collection lasted for the whole swarm duration (about 20-25 min). We chose a swarm far from habitations and livestock to limit odor pollutions, containing a large number of males (more than 1,000) and attractive for females (observation of a large number of couples). After VOC collections, twisters were individually placed into glass vials and stored at 4 °C until analysis. Five replicates were performed on the same swarm on different days.

#### Solvent extraction

About 1,000 4- to 6-day-old males of *An. gambiae* or *An. coluzzii* were introduced into a 30 × 30 × 30 cm cage and kept under insectarium conditions. The cage was made of unpainted metallic frames covered by white nets and the bottom was covered with a white paper. Thirty minutes before sunset time, the cage was placed outside and mosquitoes were observed. A 5 × 5 cm black cloth was placed in or out of the cage according to the species as described above to stimulate swarming behavior. About 5 min after mosquitoes started to swarm, the entire cage was quickly placed into a freezer at −20° C for about 10 min. Then, mosquitoes were transferred into a 20 ml glass tube and covered with a 1:1 mix of Hexane and Dichloromethane. The tube was sealed and mosquitoes were kept in the solution for 24h at 27±2 °C. Then, the solution was filtered on glass wool to remove mosquito scales and stored at 4° C until analysis. Three different controls were made: i) with 1- to 2-day-old virgin males (young male control), ii) with 2- to 3-day-old virgin females (female control) and iii) with solvent mix alone (blank control). A total of 8 swarm extracts were done; 4 of *An. coluzzii* and 4 of *An. gambiae* and 1 replicate of each type of control per species.

#### VOC collection with SPME fiber

We replicated the experiment from the study by Mozūraitis et al. (2020) in which they collected mosquito headspace with a SPME fiber in a 1L bottle. However, because most methods used to introduce mosquitoes into a bottle may potentially result in entering pollutants as well, we added a supplementary control. One way to easily introduce the mosquitoes is to blow them from a mouth aspirator into the bottle. This classic method may bring breath volatiles within the recipient, which can contain the five compounds described in Mozūraitis et al. (2020) (Poli et al. 2010; Calenic and Amann 2014; Filipiak et al. 2014; Cainap et al. 2020). Consequently, we added a “breath control”. In addition, we also replicated the experiment with females to check for sex specificity. In the morning, the two glass bottles were cleaned with acetone and dried under a clean air flow, then closed with nalophan. Three hours before swarming time (5pm), we collected the headspace of empty bottles with polydimethylsiloxane/divinylbenzene-coated SPME fibers for 1h (step 1: empty bottle). Then, we simulated the introduction of mosquitoes by blowing 3 times for 3 seconds with a mouth aspirator into the bottles. These were then closed with Nalophan (6pm) and the head space of the breath of the experimenter was collected for 1h with SPME fibers (step 2: breath control). We then introduced 60-70, 5- to 7-day-old unmated males in each bottle with the mouth aspirator, closed the bottles with Nalophan (7pm) and let the mosquitoes acclimate for 1h. During the acclimation period, we simulated a light transition from photophase to scotophase to trigger crepuscular flight. Then, when males started to fly (8pm), SPME fibers were introduced for 1h (step 3: “swarm” extract). Six different fibers were used and cleaned at 250 °C during 5 min in the injector of a gas-chromatograph (HP 6890 Series PLUS, Agilent, Santa Clara, USA) just before use. Each fiber was always used for the same type of extract and in the same bottle. Before the experiment, we made sure that the fibers had similar sensitivity by exposing them to a mix of the five compounds. Only minor quantitative differences were detected, and this was considered during subsequent statistics. Secondly, we performed the same 3 steps but with 60-70, 3- to 4-day-old virgin females instead of males. And finally, as during the introduction of mosquitoes, we blew a second time into the bottles and repeated the three steps of headspace collection but this time, instead of introducing mosquitoes in the 3^rd^ step, we blew again into the bottles (breath ×2) to check for potential accumulation of breath compounds. We kept track of the day of the experiment, the bottles and fibers identity for statistical analyses.

### Chemical analyzes

Samples obtained on Porapak-Q VCT tubes were eluted with 150 μl of Hexane and injected (1 μl) into a Gas Chromatograph coupled with Mass Spectrometer (GC-MS). Those obtained on Tenax/Carbotrap tubes, twisters or SPME fibers were directly thermo-desorbed into GC-MS. Solvent extracts were injected as is or concentrated under a gentle stream of nitrogen. All the analyses were run at the “Platform for Chemical Analyses in Ecology” (PACE), the technical facilities of the LabEx CeMEB (Centre Méditerranéen pour l’Environnement et la Biodiversité, Montpellier, France).

#### Liquid analysis (Porapak elutions and solvent extractions)

Liquids were analysed with a GC-MS QP2010 Plus (Shimadzu, Kyoto, Japan). Mass spectra were recorded in electronic impact mode (EI) at 70eV over a m/z mass range from 38 to 350. The temperature of the transfer line and the ion source were programmed to 250 °C and 200 °C, respectively. The injections were made with an injector temperature of 250 °C, and a 1:4 split mode ratio. Analyses were performed using a 30 m × 0.25 mm × 0.25 μm Optima 5-MS (Macherey-Nagel, Düren, Germany) fused silica capillary column with a constant helium flow set close to 1 ml.min^-1^. The oven temperature was programmed as follows: 40 °C (held 5 min) to 250 °C at 6 °C.min^-1^, and finally to 300 °C at 14 °C.min^-1^ and held 2 min. GC-MS Solution software (Shimadzu, Kyoto, Japan) was used for data processing of these analyses, with the NIST 2011 as spectrum database.

#### Tenax/Carbotrap tubes and twister analysis

These sorbents were analysed as described by Souto-Vilarós et al. (2018). Samples were analysed using a gas chromatograph (GC, Trace™ 1310, Thermo Scientific™ Milan, Italy) coupled with a mass spectrometer (MS, ISQ™ QD Single Quadrupole, Thermo Scientific™ Milan, Italy). The column used was an Optima 5-MS, the same as for liquid analysis. Absorbent traps were handled with a Multi Purpose Sampler (Gerstell, Mülheim, Germany) and desorbed with a double stage desorption system, composed of a Thermal Desorption Unit (TDU) and a Cold Injection System (CIS) (Gerstell, Mülheim, Germany). The sorbents were desorbed in splitless mode at 250 °C on a trap cooled at −80 °C by liquid nitrogen. Then, the trap was heated to 250 °C with a 1:4 split ratio to inject the compounds in the column. The carrier gas used was helium at 1 ml.min^-1^. Oven temperature program was as follows: held at 40 °C for 3 min., then increased to 220 °C at 5 °C.min^-1^ and finally to 250 °C at 10 °C.min^-1^, and held for 2 min. Mass spectra were recorded in electronic impact mode (EI) at 70eV over a m/z mass range from 38 to 350. The temperature of the transfer line and the ion source were programmed to 250 °C and 200 °C, respectively. Xcalibur™ software (Thermo Scientific™, Milan, Italy) was used for data processing and the NIST 2011 as spectrum database.

#### SPME fiber analysis

We analyzed SPME fibers as described in Mozūraitis et al. (2020) with a GC-MS QP2010-SE (Shimadzu, Kyoto, Japan). Volatiles were desorbed into the injector at 225 °C for 1 min in split mode (1:4 ratio). Analyses were performed using a 30 m × 0.25 mm × 0.25 μm J&W DB-Wax silica capillary column (Agilent, Santa Clara, USA) with a constant helium flow at 1 ml.min^-1^. The oven temperature was programmed as follows: 40 °C (held 1 min.) to 150 °C at 5 °C.min^-1^, and finally to 220 °C at 20 °C.min^-1^ and held 9 min. Mass spectra were recorded in electronic impact mode (EI) at 70eV over a m/z mass range from 38 to 350. The temperature of the transfer line and the ion source were 250 °C and 200 °C, respectively. GC-MS Solution software (Shimadzu, Kyoto, Japan) was used for data processing of these analyses.

### Electrophysiological analysis

Electrophysiological activity of female olfactive detection, on the male swarm extracts, was tested with an Electro-Antennografic Detection system, coupled with Gas Chromatography (GC-EAD) at the PACE. Thirty minutes before the mosquito scotophase, both 4- to 6-day-old males and 2- to 3-day-old virgin females were placed separately in cardboard cups (Ø = 75 mm, h = 100 mm), provided with a 5% honey solution and transferred from the insectarium to the electrophysiology laboratory at PACE. At mating time, mosquitoes were placed in the dark. Males were checked for erected antennae as a proxy to ensure that change in the light-dark cycle, transport and electrophysiology laboratory conditions did not alter mosquito motivation for mating. Two minutes after male antennae were erected, a female was removed from the cardboard for the antennal preparation.

The female’s head was gently excised with a pair of surgical scissors and mounted between two glass capillary tubes of 76 mm in length and 1.12 mm in diameter (World Precision Instrument, Sarasota, USA), pulled and cut using a vertical micropipette-puller (P-30 model, World Precision Instruments, Hertfordshire, United Kingdom) filled with insect Ringer solution (6.0 g.l^-1^ NaCl, 0.4 g.l^-1^ KCl, 0.27 g.l^-1^ CaCl_2_ and 3.20 g.l^-1^ NaC_3_H_5_O_3_) and connected to silver wires. The indifferent electrode was inserted into the back (the foramen) of the isolated head and the intact tip of one antenna was inserted into the recording electrode which was connected to an Electro-Antennography Detector setup (EAD, Syntech IDAC-2, Kirchzarten, Germany). One end of the glass electrodes was pulled to a fine point which was then broken to ensure a close fit with the female’s head and antenna.

The above EAD setup was linked to a Gas Chromatograph-Flame Ionization Detector (GC-FID, CP-3800, Varian, Palo Alto, USA) equipped with an Optima 5-MS capillary column, the same as for the GC-MS analyses. Liquid injections after Porapack solvent elution were made in a 1079 PTV injector (Programmed Temperature Vaporizator) at 250 °C with a 1:4 split ratio. Microtraps were desorbed in the same injector using a ChromatoProbe sample introduction device (Varian, Palo Alto, USA) with the same split ratio and temperature programmed as follows: increased from 40 °C to 250 °C at 200 °C.min^-1^ and held for 5 min. Oven temperature was held at 50 °C for 0.40 min, then increased to 180 °C at 10 °C.min^-1^, and finally to 220 °C at 12 °C.min^-1^, for a run duration less than 17 min. The carrier gas used was helium at 1 ml.min^-1^. The effluent was split equally into two deactivated fused silica capillary columns (100 cm x 0.25 mm), one leading to the FID (270 °C) and the other into a heated EAD port (200 °C) (transfer line: Syntech, Kirchzarten, Germany) and leaded to antenna. The GC effluent for antenna was mixed with charcoal-filtered, humidified airflow (300 ml.min^-1^). StarWorkstation 6.41 software (Varian, Palo Alto, USA) was used for data processing.

Different types of male swarm extracts were tested on the antennae of several conspecific females. Antennae of 19 *An. coluzzii*females were tested with solvent extracts, Porapak-Q VCT tube extracts and Microtrap extracts (8, 9 and 2 females, respectively) and 20 *An. gambiae* females were tested with solvent and Porapak-Q VCT tube extracts (10 females each). Before GC-EAD analysis of the VOC extracts, we checked the living antenna for good activity using a negative (solvent) and a positive (mix of many VOCs) stimulus solutions. We proceeded as follows: 5μl of each solution was applied to a 1 cm × 2 cm filter paper contained in a Pasteur pipette. The pipette was then placed in an air pump system and the volatiles were directly sent to the antennal preparation with a pulse duration of 0.5 s and a flow rate of 890 ml.min^-1^ regulated by a CS-55 Stimulus Controller (Syntech, Hilversum, Netherlands).

### Identification of specific and active VOCs

For all the chemical analysis, we compared the chromatograms of swarming mosquito extracts with those of their corresponding controls to search for any qualitative or quantitative differences. When such a difference was pinpointed, we matched the mass spectra of the compounds of interest with the NIST 2011 MS library. A solution of n-alkanes (Alkanes standard solution, 04070, Sigma Aldrich, Darmstadt, Germany) was also injected to calculate the linear retention index (LRI) of these compounds of interest (Zellner et al. 2008) and their LRIs were compared with those reported in the literature.

In the electrophysiological analysis, we studied the chromatograms in a range of LRI between 800 and 1700, including compounds present at very low and trace amounts. When a depolarization was observed in the electrophysiological analysis, a section of 20 points of LRI was studied around the compound’s LRI on the GC-MS trace. Comparison of the samples with their respective controls collected on the same day, allowed to subtract potential contaminants from the samples. Non-natural compounds, such as industrial chemicals and/or compounds not naturally produced by living organisms (Charpentier et al. 2012), were considered as contaminants.

We also particularly searched for the presence of acetoin, sulcatone, octanal, nonanal and decanal reported by Mozūraitis et al. (2020). With that aim, we defined a LRI window of 20 points on both sides of the LRI of these compounds on the bases of injected standards (for SPME fibers; acetoin, sulcatone, octanal, nonanal and decanal, ≥98%, 99%, 99%, ≥98% and ≥98% GC, respectively, Sigma-Aldrich, Darmstadt, Germany) or literature (for other extraction methods). Then, we searched for their specific ions at those LRI as follows: (LRI/ion) 711/88, 981/108, 998/84, 1100/98, and 1201/112 for acetoin, sulcatone, octanal, nonanal and decanal, respectively. Then, when a peak corresponding to both LRI and specific ion was found, we compared its spectra with the NIST 2011 MS library before quantification with integration units.

### Statistical analysis

Both the activation rate and choice were analyzed using binomial Generalized Linear-Mixed Models (GLMM, “*glmer*” function in “*lme4*” package). Tested combinations (four levels: *An. coluzzii, An. gambiae, An. coluzzii vs. An. gambiae* and control tests), female species (two levels: *An. coluzzii* and *An. gambiae*) and their interaction were considered as fixed effects, and replicates as random effects. All analyses were performed using R (version 3.4.0).

We analyzed separately the quantities using integration units of acetoin, sulcatone, octanal, nonanal and decanal. When pairing between the controls and the mosquito extracts was possible (with SPME and twister) and to account for repeated measurements, we used Gaussian GLMMs. In those cases, the extract (3 levels for SPME fibers: empty bottles, breath and flying mosquitoes (either male or females or breath ×2) and 2 levels for twisters: control and swarm) was considered as fixed effect. The replicates (i.e. the day for twisters and the SPME) and fiber nested within the bottle for SPME fibers were considered random effects. Considering the SPME fibers as a random effect made it possible to account for their minor differences of sensitivity in the model. When pairing between the controls and the mosquito extracts was not possible (Tenax-TA/Carbotrap and Porapak-Q VCT tubes), Gaussian GLMs were used. The extract (empty box, *An. coluzzii* swarm or *An. gambiae* swarm) was considered as fixed effect. In some occasions, some compounds were not found in the extracts or only in two samples. Consequently, it was not possible to perform statistical analyses and we considered the extracts as not significantly different. No statistical analysis was performed with liquid extracts due to a small number of replicates. For model selection, we used the stepwise removal of terms, followed by likelihood ratio tests. Term removals that significantly reduced explanatory power (*P* <0.05) were retained in the minimal adequate model. All percentages are provided with their 95% confident interval.

## RESULTS

### Behavioral assays

Over 12 replicates of c.a. 200 females for each of the four combinations, a total of 8,918 females were tested (4,423 *An. coluzzii* and 4,495 *An. gambiae*) among which 3,591 flew upwind into the collecting boxes (activation rate: 40.2±1%). There was a significant interaction between female species and the four combinations of swarm boxes tested (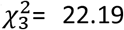, *P* <0.001). Species subset analyses showed that the different test combinations had no effect on the *An. coluzzii* activation rate (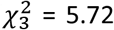, *P*=0.12; Fig. 2). However, in *An. gambiae,* there was a significant effect of the test combinations on the female activation rate (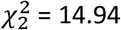, *P* = 0.001; Fig. 2) with a higher activation rate when females were exposed simultaneously to both *An. coluzzii* and *An. gambiae* male swarms than when exposed to an *An. coluzzii* swarm alone (*z* = 4.01, *P* <0.001) or to the control (two empty boxes: *z* = 2.57, *P* = 0.04). Nonetheless, no difference was found when females were exposed to an *An. gambiae* male swarm alone (*z* = −1.49, *P* = 0.44). Neither the *An. coluzzii* swarms, nor the *An. gambiae* swarms were attractive for activated females (female species: 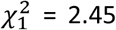, *P* = 0.11; choice test combinations: 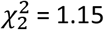, *P* = 0.56 and female species × choice test combination interaction: 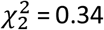, *P* = 0.84; Fig. 3).

**Fig. 2:**
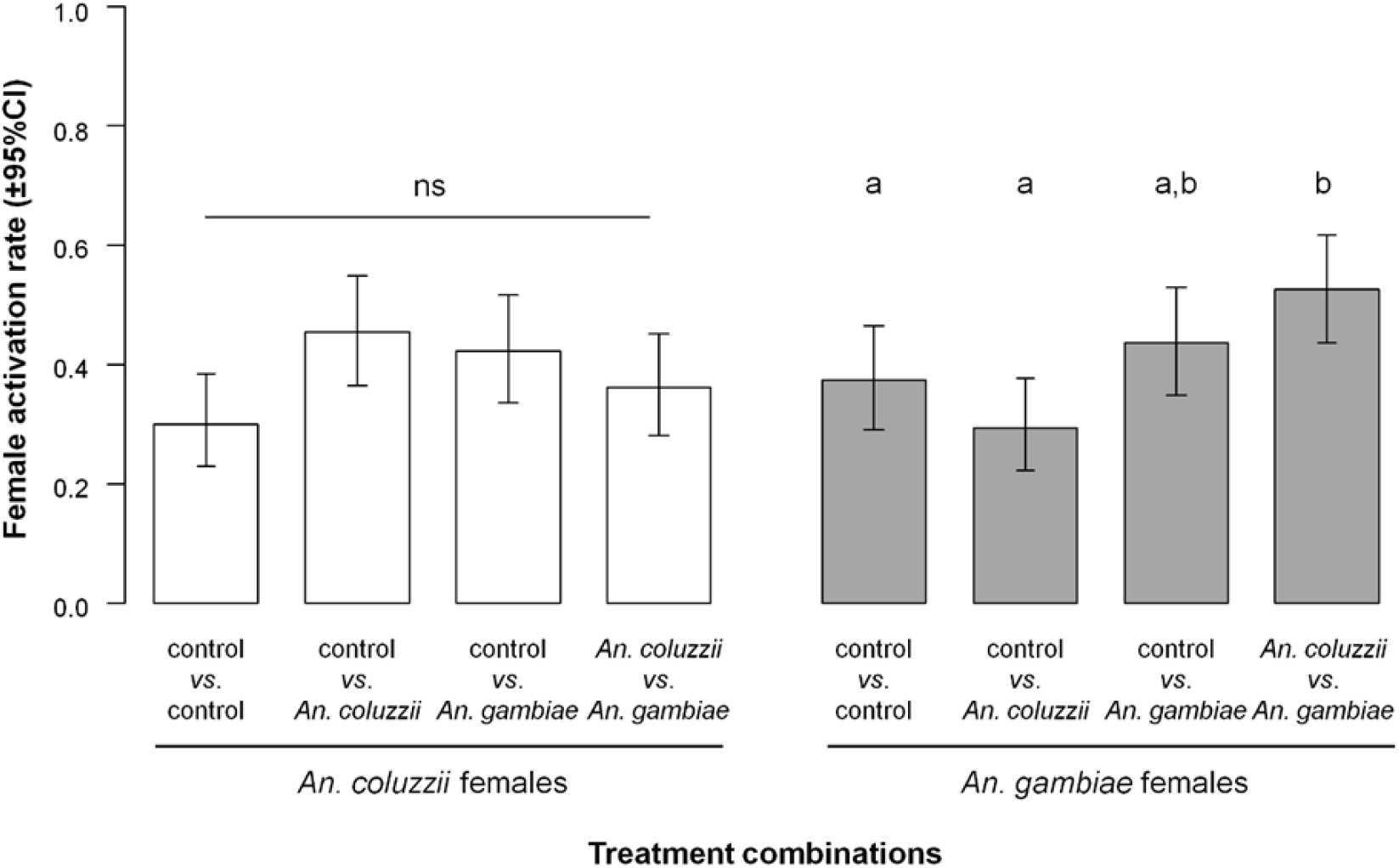
Mosquito activation rate, expressed as the proportion of females caught in both collecting boxes out of the total number released females for each of the four tested combinations. Different letters indicate difference at *P*<0.05.

**Fig. 3:**
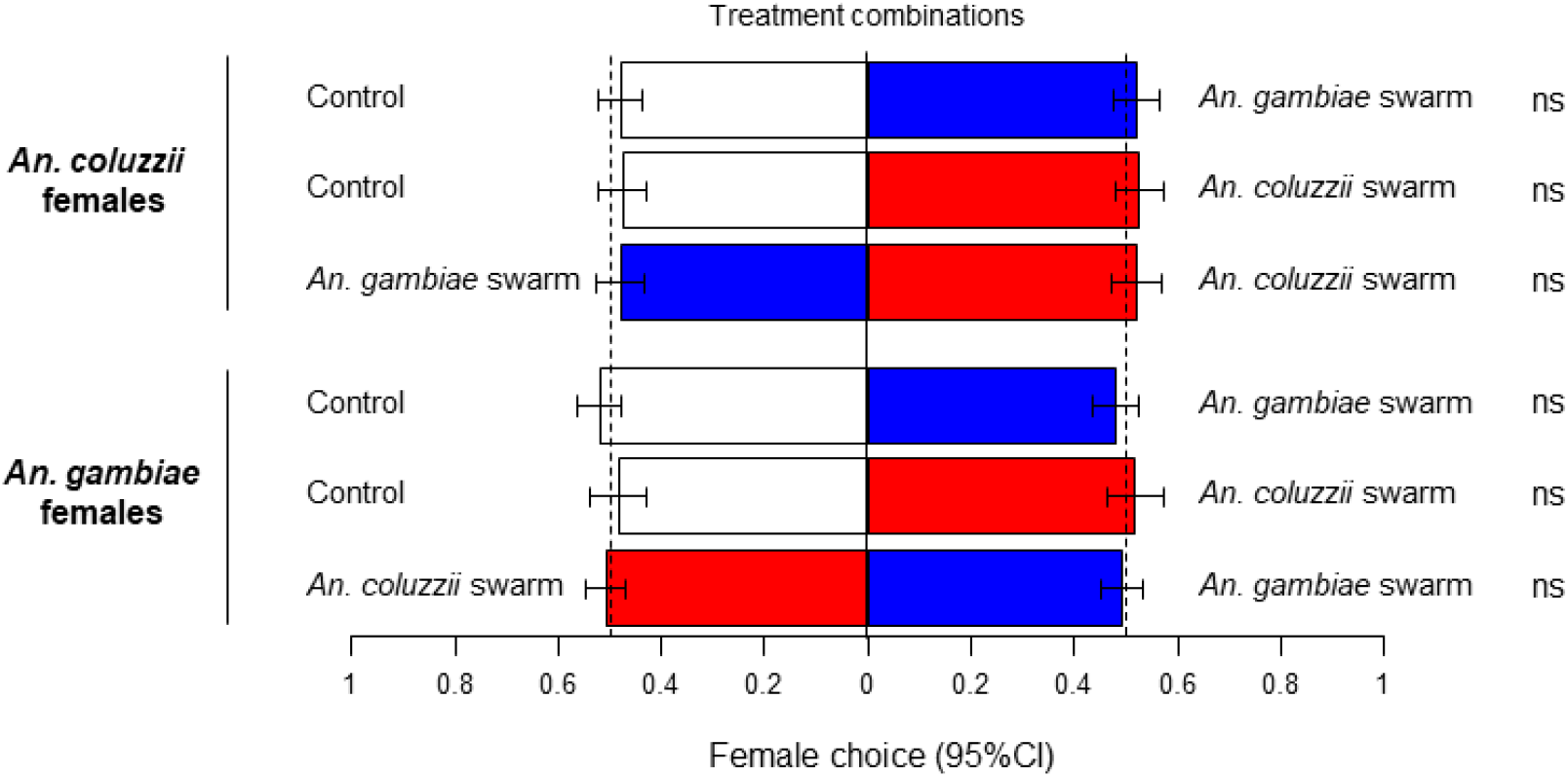
Mosquito choice, expressed as the proportion of females caught in one or the other collecting box out of the total number of “activated” females for each test.

### Chemical analyses

No qualitative or quantitative differences were detected between samples of swarming males and their relative controls for both species obtained from the Porapak-Q, Twisters® and the liquid extracts (Figs S2 & S3). Minor quantitative differences were detected with the Tenax/Carbotrap tubes. However, mass spectra showed that these peaks were pollutants, such as silicones, BTX (mono-aromatic compounds like benzene, toluene and xylene), alkanes which were all emitted by the box containing the mosquitoes. Quantitative differences including linear organic acids were also detected, but they were not reproducible.

During the experiment replicating the protocol by Mozūraitis et al. (2020), both males and females started to fly at the expected time (8pm) but due to the small volume it was not possible to determine if the group was a swarm. Analyses of SPME fibers in search for acetoin, sulcatone, octanal, nonanal and decanal showed that acetoin was absent from almost all the male samples (Fig. 4A), but was present on three occurrences in female extracts (over six) (Fig. 4B). Sulcatone, octanal, nonanal and decanal were found in almost all types of extracts including controls. However, we did not find significant increased quantities of the five compounds in “swarming” mosquito extracts compared to the breath controls (Fig. 4). The control to test the effect of blowing twice in the bottle showed that the headspaces of “breath” and “breath ×2” were very similar (Fig. S4), demonstrating an absence of accumulation of breath odor between the breath control and the introduction of mosquitoes.

**Fig. 4:**
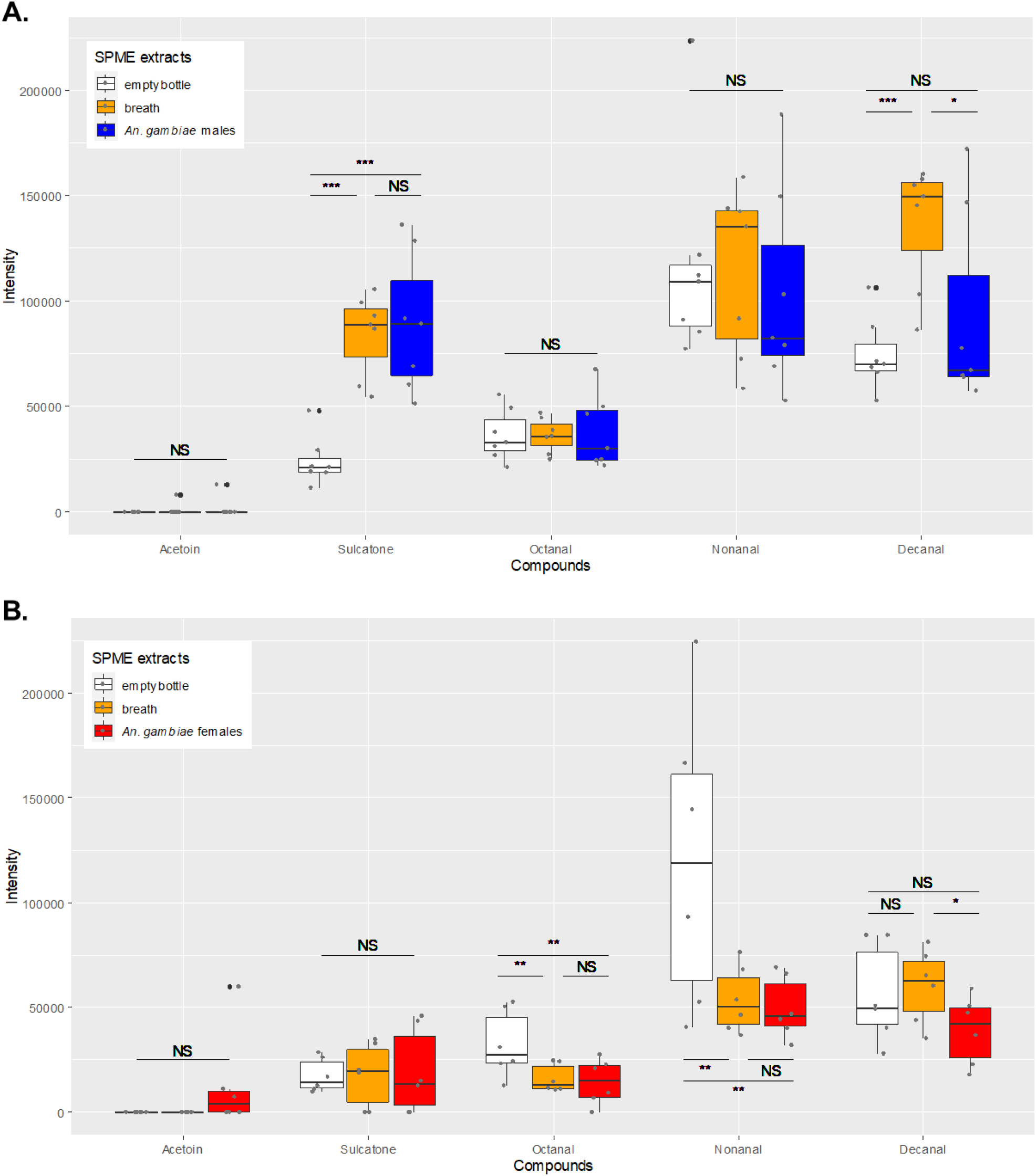
Quantities (integration units) of acetoin, sulcatone, octanal, nonanal and decanal contained in the headspace of the empty bottles, the breath of the manipulator and flying males (A) and females (B) of *Anopheles gambiae* Kisumu and collected with SPME fibers. The intensity values correspond to the counts related to the abundance of the specific ions representative of each molecule formed in the mass spectrometer and correspond to the amount of compound analyzed. The box plots indicate the median (wide horizontal bars), the 25th and 75th percentiles (squares), and the minimum and maximum values (whiskers). The black dots represent outliers and grey dots raw data. NS=not-significant; * =P < 0.05; ** =P < 0.01; *** =P < 0.001.

A posteriori, we also quantified acetoin, sulcatone, octanal, nonanal and decanal in our previous GC-MS analyses in search for similar variations as described in Mozūraitis et al. (2020). We did not find acetoin in our chromatograms. Indeed, as we used a low polarity Optima 5-MS column, acetoin (if present) was eluted before the LRI 800. In the samples obtained from natural swarms of *An. coluzzii* with the twister method, sulcatone and octanal were absent. The quantities of both nonanal and decanal were not significantly higher in the swarm than in the control (Fig. 5A). In laboratory swarm samples, sulcatone, octanal, nonanal and decanal were found inconsistently. Nevertheless, swarms did not show higher quantities of these compounds compared to the empty box (Fig. 5B) excepted in samples obtained with the Porapack tubes (Fig. 5C). However, this difference was not statistically significant. Whatever, the method used, we can note a high quantitative variability between the samples. This was true for both the control and the mosquito samples.

**Fig. 5:**
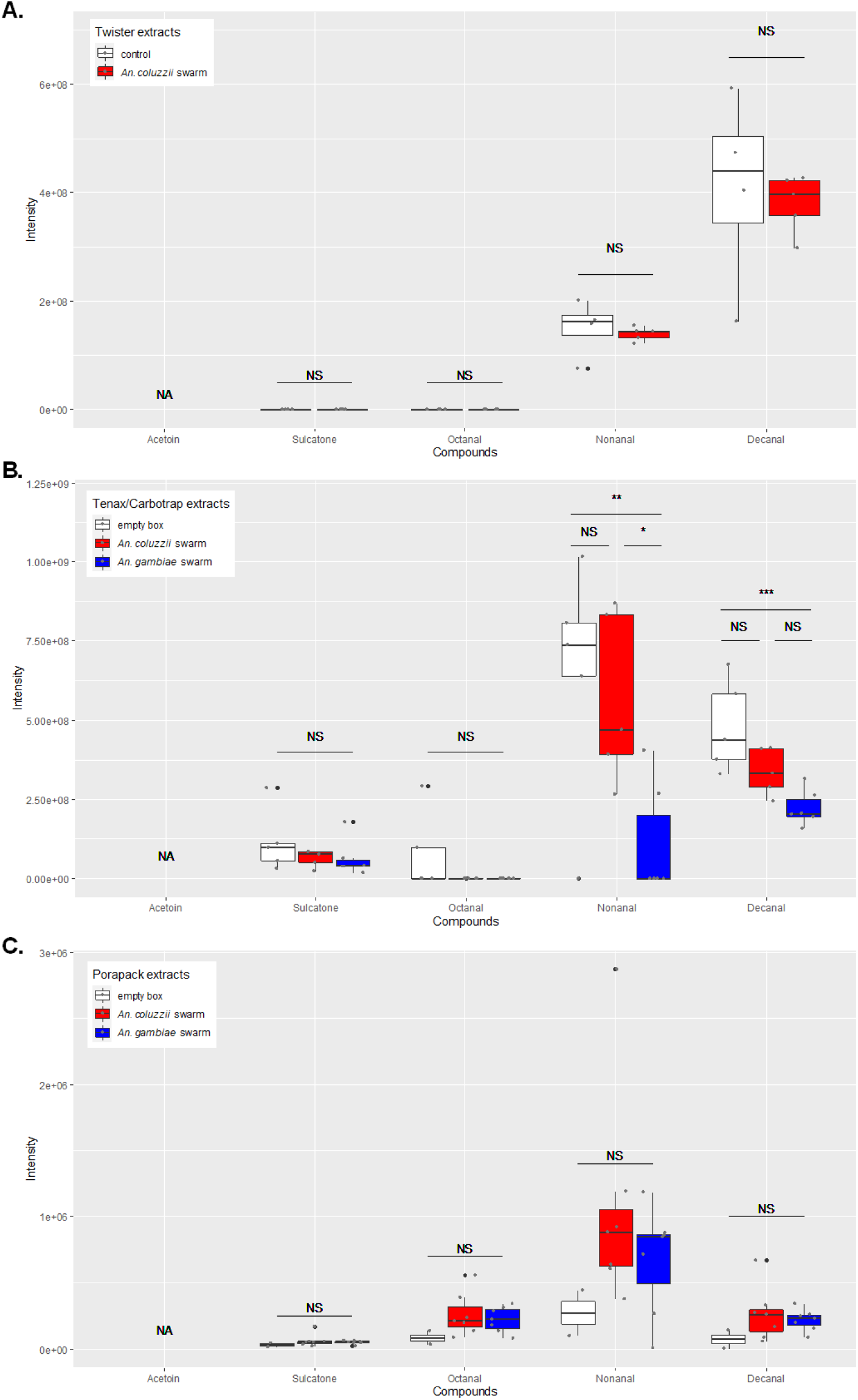
Quantities (integration units) of sulcatone, octanal, nonanal and decanal contained in a natural swarm sampled with twisters (A); in the headspace of the empty box, swarming *Anopheles coluzzii* males and *An. gambiae* males sampled with Tenax/Carbotrap tubes (B); or collected with Porapack tubes (C). The intensity values correspond to the counts related to the abundance of the specific ions representative of each molecule formed in the mass spectrometer and correspond to the amount of compound analyzed. The box plots indicate the median (wide horizontal bars), the 25th and 75th percentiles (squares), and the minimum and maximum values (whiskers). The black dots represent outliers and grey dots raw data. NS=not-significant; * =P < 0.05; ** =P < 0.01; *** =P < 0.001. NA: The analytical procedure for these analyses did not allow detection of acetoin.

## DISCUSSION

For the past decades, several authors have suggested the existence of long-range sex pheromones in the attraction of *An. gambiae s.l.* females to male swarms (Charlwood et al. 2002; Tripet et al. 2004; Diabaté et al. 2011; Poda et al. 2019) but to date, no reliable experimental evidence is available. Here, we used behavioral, physiological and detailed chemical approaches to attempt to highlight the existence of such compounds. However, despite a large set of methodologies and working hypotheses to maximize the chances of highlighting the presence of pheromones (*i.e.* timing, physiological state, natural behavior, recently colonized mosquitoes), our study failed to provide evidence of the presence of long-range male sex pheromones in both *An. coluzzii* and *An. gambiae.* We also replicated an experimental setup that recently showed an increase in quantity of five compounds in “swarming” males compared to controls. However, unlike in Mozūraitis et al. (2020), we did not find any significant difference with our controls.

During our behavioral tests in the olfactometer, females were not attracted to air currents passed over male swarms regardless of the species. During the experiment, the males released in the large plexiglass boxes and exposed to natural light started to fly randomly at sunset. Then, about 150-200 males out of the 500 released (<50%) gathered in the upper half of the box flying in loops without touching the box walls. They also reacted to the movements of the visual marker, meaning they were indeed swarms. The other males flew randomly in the box, bouncing against the walls. Females showed a good activation rate but they did not show a particular choice for any arm, containing a swarm or not. Nevertheless, *An. gambiae* females showed an intriguing higher activation rate when exposed simultaneously to both a conspecific and a heterospecific swarm compared to a single *An. coluzzii* swarm. This was, however, probably without biological significance as no difference was detected in the test providing *An. gambiae* females with a choice between the control and *An. gambiae* swarm, *vs.* a test offering two controls.

Our behavioral result was consistent with our physiological study. Indeed, despite the fact that we checked for both behavioral receptivity of mosquitoes and for receptivity of mounted antennae, no consistent antennal depolarization was observed in females across assays when exposed to swarm extracts. This suggests a lack of response by females to any volatile chemicals present in our swarm extracts.

Chemical analyses were also negative and no compounds specific for male swarms could be detected whatever the method used. Moreover, unlike in Mozūraitis et al. (2020), the quantities of acetoin, sulcatone, octanal, nonanal and decanal were found inconsistently across swarming mosquito samples and not in larger quantities compared to controls, making it difficult to support the assumption according to which they could actually be emitted by males. These divergent results could be explained by the fact that, unlike in the experiments of Mozūraitis et al. (2020), we used a control which considered the potential introduction of pollutions at the same time as mosquitoes. This showed that breath was responsible for most of the variability. The most convincing result showing that sulcatone, octanal, nonanal and decanal are probably not pheromones emitted by males is the one obtained with the twister method. Indeed, twisters are coated with the same adsorbent phases as SPME fibers used both in this study and in the one by Mozūraitis et al. (2020). According to the manufacturer, twisters are up to a thousand times more sensitive than SPME fibers partly due to a larger sorbent volume. Moreover, we exposed the twisters directly in a natural swarm formed by more than 1 000 males (probably up to 6 000 males according to the estimation of trained technical staffs).

Despite this sensitivity, the number of mosquitoes and the natural biological context, sulcatone and octanal were not found and both nonanal and decanal had similar quantities in the swarm and outside the swarm (control twister placed 3 m upwind from the swarm). In addition, the high quantitative variability found in the laboratory and the fact that these compounds are frequently found in controls suggest that they could be laboratory and/or human pollutions that are difficult to control for.

Our results contrast with those of Mozūraitis et al. (2020) probably for several reasons. First, in their laboratory experiments, they reported a simple flight activity instead of a swarming activity in which males should fly in regular loops with erected antennae (Downes 1969; Poda et al. 2019). This can be explained by the absence of adequate visual stimuli such as ground markers which are mandatory to trigger swarming behavior (Charlwood and Jones 1980; Marchand 1984; Gibson 1985; Facchinelli et al. 2015; Niang et al. 2019; Poda et al. 2019). Second, this flight activity was extraordinary long for a swarming flight, exceeding 200 min in the presence of the five compounds. Indeed, swarming behavior is known to last for only 20-30 min in nature (Charlwood and Jones 1980; Marchand 1984; Sawadogo et al. 2013, 2014; Bimbilé Somda et al. 2018) and only up to 60 min in artificial conditions (Charlwood and Jones 1980; Poda et al. 2019). In the latter, the number of males in swarms decreased over time while the others flew randomly, bouncing on the flight arena walls (Poda et al. 2019). Indeed, swarming is an activity with a high energy demand, consuming 50% of sugar and glycogen reserves in 25 min (Maïga et al. 2012, 2014). Mosquitoes probably switched to a more random flight to try to escape and search for a sugar meal to refuel their reserves. This behavior can be stimulated by acetoin, sulcatone, octanal, nonanal and decanal, as they are frequently emitted from nectar sources and fermented sugar (Goodrich et al. 2006; Schiestl 2010; Dekel et al. 2019). Finally, they also reported that these five compounds increased insemination rates in five different species in semi-field cages. However, instead of stopping the experiment after the swarming time, they collected the females in the morning. As they left both a dark box in the arena as resting site and a sugar source, it is likely that males and females also mated in or around these two resources during the night. Indeed, it was shown that mating can occur indoors, outside of swarming times and locations (Dao et al. 2008). The boxes and sugar sources with these five compounds were probably more attractive than the control ones gathering more mosquitoes and thus inducing more inseminations.

Independently, our methods can also have weaknesses. Even if the number of swarming males was biologically relevant compared to natural swarms, a higher number of males could be needed in experimental setups, or the sensitivity thresholds of our chemical and electrophysiological apparatus are below the ability of insects to detect low amounts of compounds. In addition, although the air flow used in our VOC collection in laboratory was not unconventional, it might have been too high inducing a leak of compounds of interest. Likewise, the air quality could have masked some compounds emitted in minute quantities even though our behavioral experiment and VOC collection in natural swarms led to negative results too.

However, altogether and keeping in mind that an absence of evidence is not an evidence of absence, our results support the absence of long-range sex pheromones emitted by male swarms. However, further investigations are needed and complementary methodologies such as electro-physiology on palps (Lu et al. 2007; Iatrou and Biessmann 2008; Pitts et al. 2011; Guidobaldi et al. 2014), molecular analyses of olfactory protein expression on adequate physiological status, or real-time chemical analyses of swarm volatiles with a sensitive apparatus such as a proton-transfer-reaction mass spectrometry could be options. The eventuality that organic compounds with low volatility might be involved in mating process other than long-range attraction cannot be discarded.

The question of how *Anopheles* females seek male swarms is still open. The lack of post-mating barrier (Persiani et al. 1986; Diabaté et al. 2005, 2007; Hahn et al. 2012; Pombi et al. 2017) and the low rate of hybrids in this geographical area (della Torre et al. 2001, 2005; Tripet et al. 2001; Diabaté et al. 2006) necessarily involve a reasonably distant process which prevents females from entering heterospecific swarms. As acoustic cues have shown some limitations at long range (Feugère et al. 2021), and because long-range chemical cues are still disputable, visual cues such as ground markers (Diabaté et al. 2009; Sawadogo et al. 2014; Poda et al. 2019) could be good candidates, but their effective range and level of specificity is poorly known.

## DECLARATIONS

### Ethical approval

The use of rabbit blood was approved by the Ministry of Higher Education, Research and Innovation under the registration No. APAFIS#13817-2018022712203932 v4.

### Availability of data and material

The raw datasets are available online: https://doi.org/10.5281/zenodo.4719568.

### Code availability

Script and codes are available online: https://doi.org/10.5281/zenodo.4719568.

### Authors’ contributions

O.R. conceived the study. O.R., S.B.P., B.B., B.L. and L.D. designed the chemical and electrophysiological experiments. O.R. and S.B.P. performed chemical extractions and SBP and BB performed the chemical analysis. S.B.P. and B.L. performed the electrophysiological experiments. O.R. and S.B.P. designed olfactometric experiments and S.B.P. performed data collection. S.B.P. and O.R. performed statistical analyses. S.B.P., O.R., B.B., B.L. drafted the manuscript and L.D., O.G., A.D, and R.K.D critically revised the manuscript. All authors revised the manuscript, gave final approval for publication and are accountable for the work performed therein.

## Acknowledgements

We thank David Sanou for his assistance in the field, Stephane Somda, Sofan Somé, Bethsabee Scheid, Marie Rossignol and Carole Ginibre for their help in mosquito rearing and Heidi Lançon for proofreading the paper. Version 6 of this preprint has been peer-reviewed and recommended by Peer Community In Ecology (https://doi.org/10.24072/pci.ecology.100091).

## SUPPLEMENTARY FILES

**Fig. S1:**
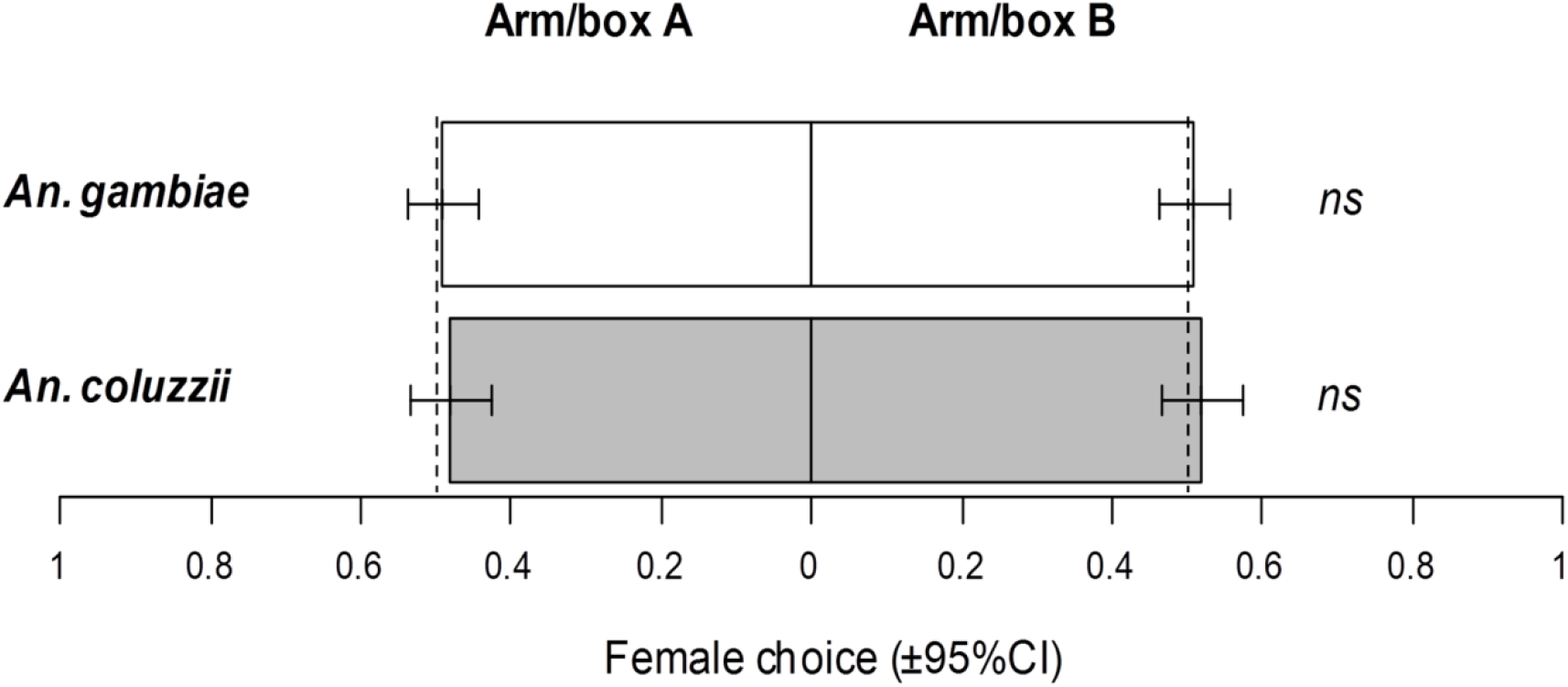
Test for olfactometer symmetry. Female choice, expressed as the proportion of female mosquitoes caught in one or the other collecting box out of the total number retrieved from both collecting boxes for the test with empty boxes (control *vs.* control combination).

**Fig. S2 A:**
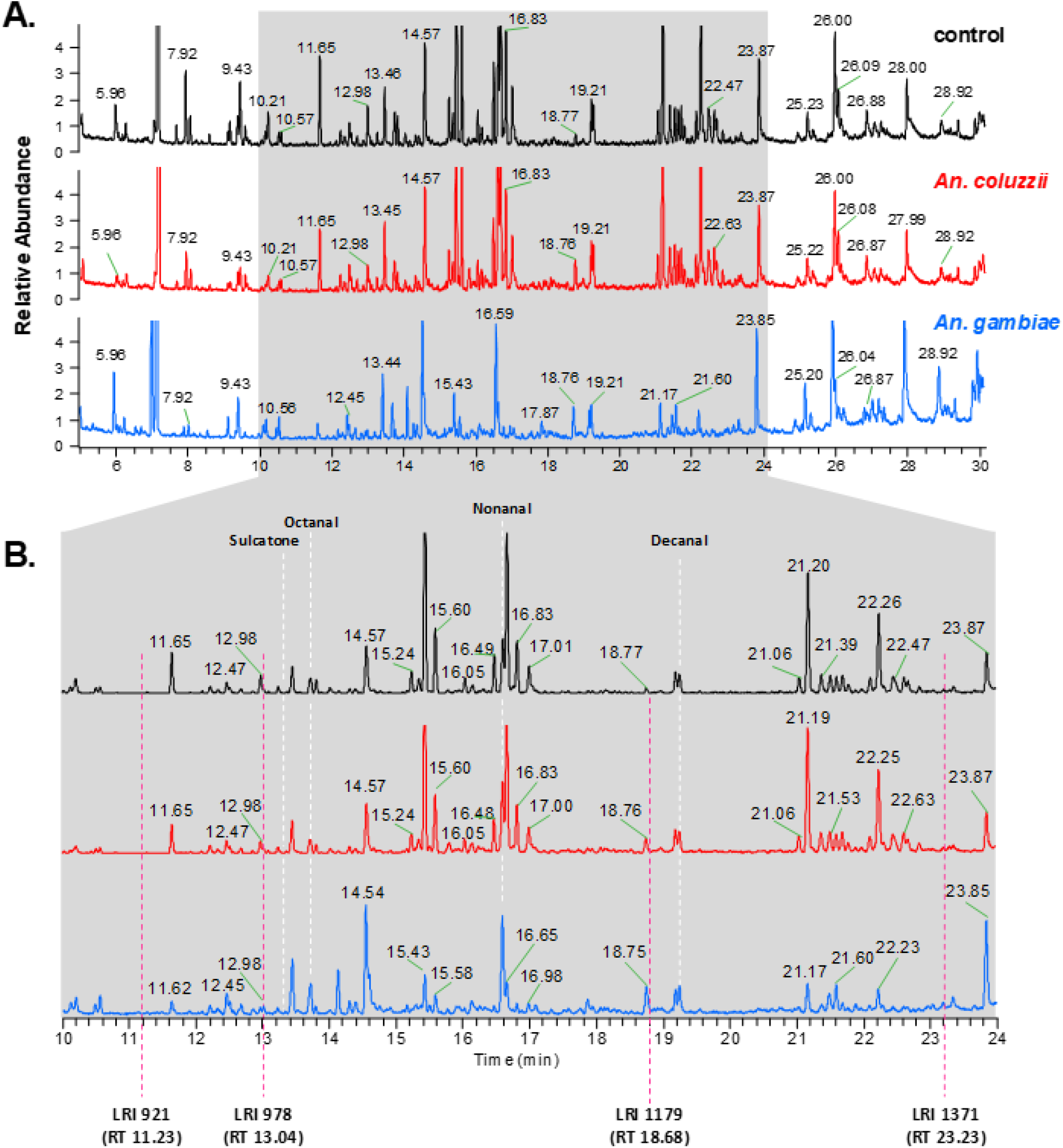
Chromatograms obtained with eluted Porapak extracts from headspaces of the control, an *An. coluzzii* swarm and an *An. gambiae* swarm (linear retention index (LRI) 800 to 1700). **B.** Enlarged view of the retention time (RT) interval containing the LRI at which *An. coluzzii* antennal responses were detected in GC-EAD. Doted white lines correspond to the theoretical RT of Sulcatone, octanal, nonanal and decanal. No difference was detected between the control and the swarm extracts. All the peaks correspond to emissions born from the material.

**Fig. S3 A:**
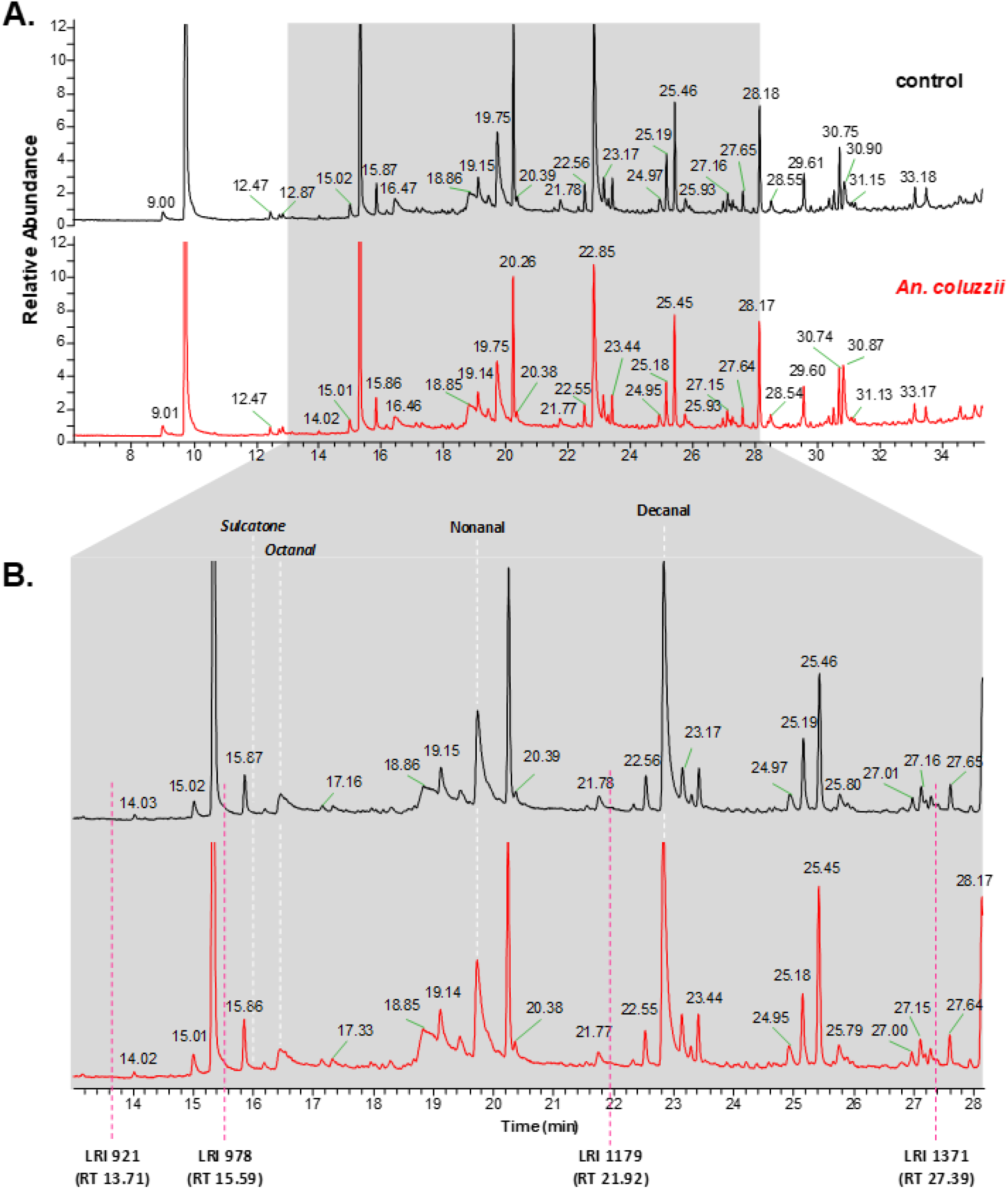
Chromatograms obtained with the desorption of twisters obtained with the static sorptive collection methods under natural conditions in the field (linear retention index (LRI) 800 to 1700). The twister was either located within a non-enclosed swarm of *An. coluzzii* or 3 m away upwind (control). **B.** Enlarged view of the retention time (RT) interval containing the LRI at which *An. coluzzii* antennal responses were detected in GC-EAD. Doted white lines correspond to the theoretical RT of sulcatone, octanal, nonanal and decanal. When in italic, the compound was absent. No difference was detected between the control and the swarm extracts. All the peaks correspond to environmental emissions.

**Fig. S4:**
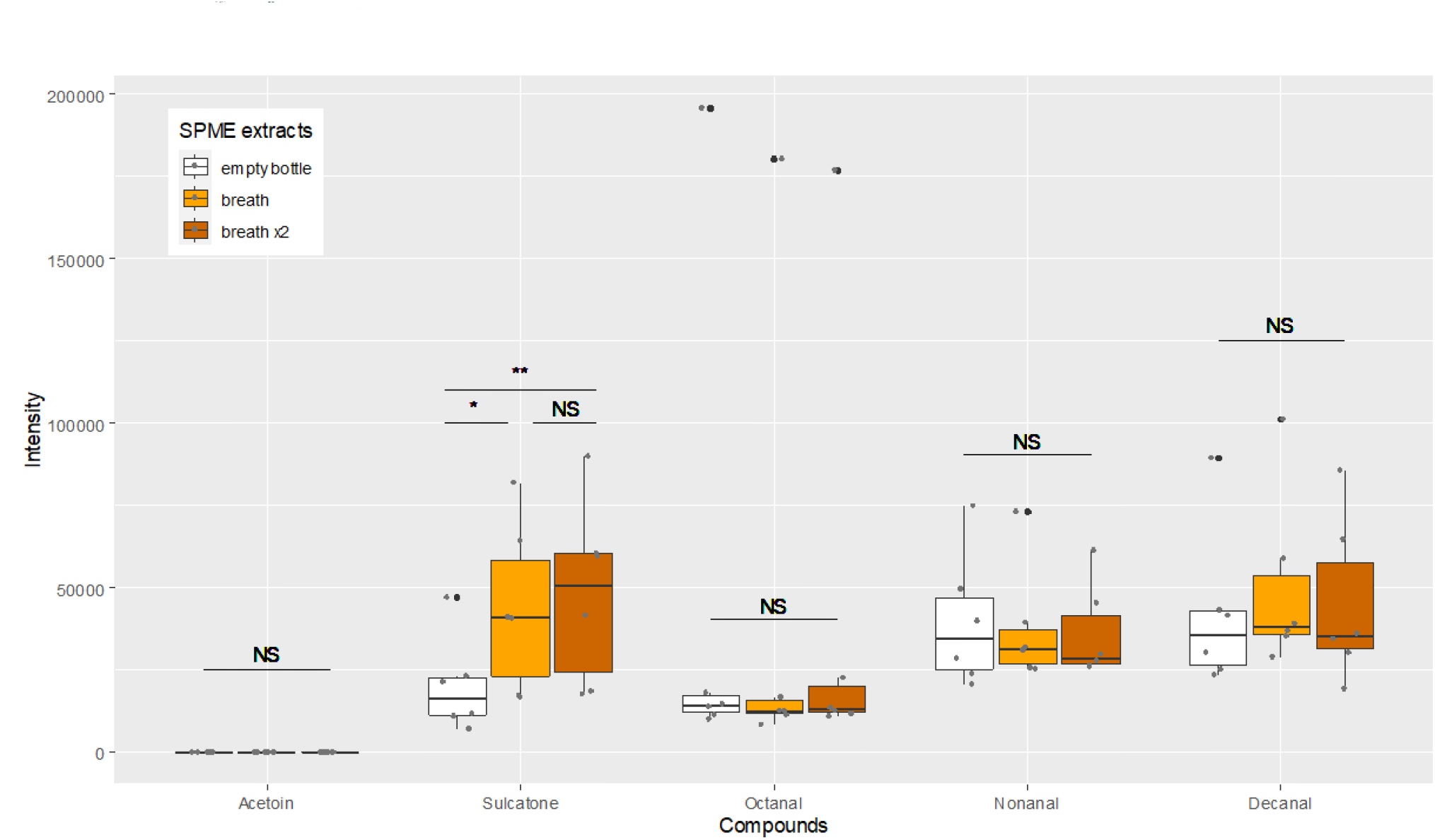
Quantities (integration units) of acetoin, sulcatone, octanal, nonanal and decanal contained in the headspace of the empty bottles, the breath of the manipulator and the breath of the manipulator after blowing for the second time (breath ×2) sampled with SPME fibers. The intensity values correspond to the counts related to the abundance of the specific ions representative of each molecule formed in the mass spectrometer and correspond to the amount of compound analyzed. The box plots indicate the median (wide horizontal bars), the 25th and 75th percentiles (squares), and the minimum and maximum values (whiskers). The black dots represent outliers and grey dots raw data. NS=not-significant; * =P < 0.05; ** =P < 0.01.

